# Homeostatic mitophagy scales mitochondrial networks

**DOI:** 10.1101/2023.12.17.572060

**Authors:** R.L. Thomas, B.C. Bishop, L.M. Andre, H. Nolte, M. Graef

**Affiliations:** Max Planck Research Group “Autophagy and Cellular Ageing”, Max Planck Institute for Biology of Ageing, 50931 Cologne, Germany; Department of Molecular Biology and Genetics, Cornell University, Ithaca 14853, NY, USA; Department of “Mitochondrial Proteostasis”, Max Planck Institute for Biology of Ageing, 50931 Cologne, Germany

## Abstract

Cells scale organelle sizes to ensure physiological function. Here, we uncover a dual role for autophagy in controlling mitochondrial network size with a key function for homeostatic mitophagy. During starvation, non-selective autophagy sustains mitochondrial biogenesis in non-dividing yeast cells resulting in mitochondrial network expansion. Strikingly, Atg32/Bcl2L13-mediated mitophagy scales mitochondria back to pre-starvation size. Without mitophagy, mitochondria size increases two-fold while retaining wildtype-like structural, compositional, and metabolic features. In turn, synthetically elevated mitophagy only mildly reduces mitochondria size, suggesting cells maintain a minimal network size by compensatory biogenesis. Single-cell analysis predicts two metabolically tunable setpoints, mitochondria-to-cell and mitochondria-to-cytosol volume ratios, for mitochondria scaling by autophagy-driven biogenesis and degradation, respectively. Our work reveals how cells use autophagy to scale mitochondria to metabolic and cellular size parameters.

**One Sentence Summary:** A homeostatic form of mitophagy controls the network size of mitochondria during starvation.

## Main text

Autophagy is a highly conserved degradative pathway critical for cellular function and health (*1*). As a defining feature of autophagy, cells form double-membrane autophagosomes in response to diverse functional and cellular signals. During formation, autophagosomes can encapsulate a broad scope of cytoplasmic substrates, termed non-selective or bulk autophagy, or targeted specific cargos including mitochondria in a process termed selective autophagy. Receptor proteins, which are either inherently bound to or bind by ubiquitinated proteins on the surface of targeted organelles, are activated and recruit the conserved autophagy protein machinery to induce autophagosome biogenesis (*2*). Receptor-mediated selective autophagy is commonly viewed as a quality control mechanism specifically targeting dysfunctional portions of organelles for degradation to maintain functional integrity (*3*). A number of receptor system have evolved for the selective turnover of mitochondria, mitophagy, in yeast and mammals (*4*). If and how cells use non-selective or selective forms of autophagy to control organelle sizes has remained unclear. Cells actively adjust the sizes of their organelles according to metabolic or functional needs. Mitochondria form dynamic tubular networks hosting a plethora of metabolic pathways suggesting the need for adaptive size regulation (*5*). Seminal work revealed that physical size of mitochondria networks increases with the volume increase of growing buds establishing a scaling relation in dividing budding yeast (*6*). Here, we studied potential functions of autophagy in mitochondria size regulation in response to nutrient stress.

### Atg32/Bcl2L13-mediated mitophagy controls mitochondria network size during starvation

We analyzed mitochondria size dynamics in the budding yeast, *Saccharomyces cerevisiae*, in response to nitrogen starvation (hereafter starvation). To measure the physical size of tubular mitochondrial networks, we generated spinning disk confocal z-stacks of live yeast cells expressing a C-terminally GFP-tagged outer mitochondrial membrane protein Om45 (*7*). Fluorescence images of mitochondria were converted into faithful 3D reconstructions using established MitoGraph V2.0 software to quantify physical parameters of total volume and length, average width, and the number of nodes and tips of mitochondrial tubules (**Fig. 1A** **and S1A**)(*6, 8*). We measured mitochondrial volume in WT cells every 2 h over a time course of 24 h of starvation. Strikingly, mitochondrial volume significantly increased during the first 8-10 h of starvation (**Fig. 1A**), indicating continued mitochondrial biogenesis despite starvation-induced cell cycle arrest. Initial mitochondrial expansion was followed by gradual volume reduction after 10 h until reaching pre-starvation size after 24 h (**Fig. 1A**). Taken together, these data reveal dynamic changes in mitochondria network size in starving cells in which an initial phase of expansion is followed by readjustment of mitochondrial volume.

**Figure 1:**
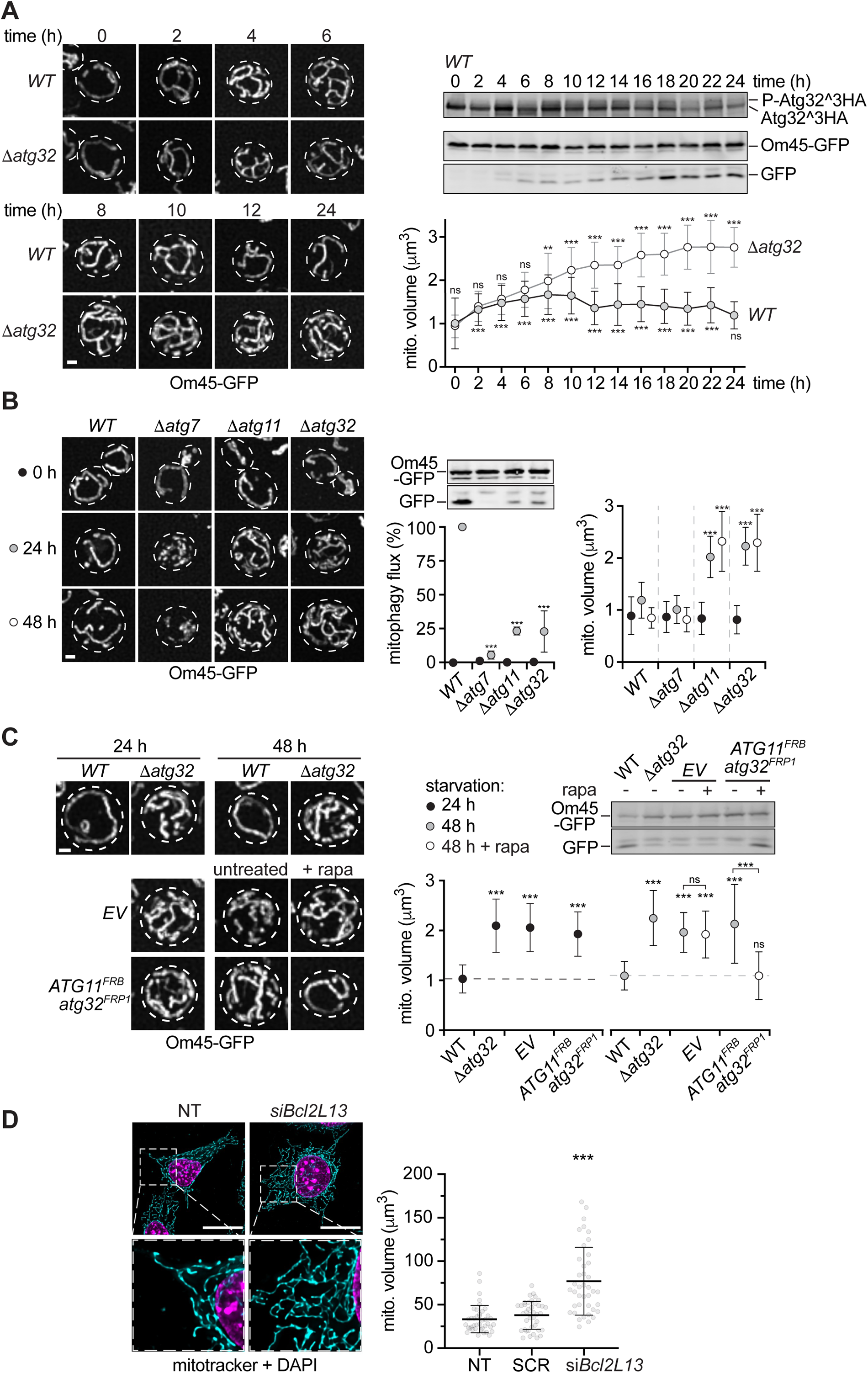
Atg32/Bcl2L13-mediated mitophagy controls mitochondrial network size during starvation. **(A)** Mitochondrial volume dynamics in WT and Δ*atg32* cells (left panels) expressing *OM45*-GFP based on fluorescence imaging (left panels) and Mitograph analysis (lower right panel) at indicated time points during starvation. Representative average intensity Z-stack projections are shown. Data are means ± SD (n=3; 60 cells/data point). Western blot analysis of WT cell lysates at indicated time points monitoring HA-tagged Atg32 phosphorylation (P-Atg32∧3HA) or Om45-GFP turnover using α-HA or α-GFP antibodies, respectively (upper right panels). Scale bar is 1 μm **(B)** Fluorescence imaging of indicated strains expressing *OM45*-GFP (left panels) and quantified mitochondrial volumes (right panel) at indicated time points during starvation. Representative average intensity Z-stack projections are shown. Data are means ± SD (n=3, 60 cells/data point). Western blot analysis of cell lysates assessing Om45-GFP turnover using an α-GFP antibody (middle panel). Data are means ± SD (n=3). Scale bar is 1 μm **(C)** Fluorescence imaging of WT or Δ*atg32* cells harboring empty vector (EV) or a plasmid encoding *ATG11^FRB^*and *atg32^FRP1^* during starvation ± rapamycin (left panel) with mitochondria volumes (lower right panel). Representative average intensity Z-stack projections are shown. Data are means ± SD (n=3, 60 cells/data point). Scale bar is 1 μm. Western blot analysis of cell lysates monitoring Om45-GFP turnover using an α-GFP antibody (upper right panel). **(D)** Fluorescence imaging of MEFs untreated (NT) or treated with scrambled (SCR) or siRNA against Bcl2L13 (siBcl2L13) after Mitotracker and DAPI staining. Mitograph-derived mitochondrial volumes are shown in right panel. Data are means ± SD (n=4, 40 cells for each condition). Scale bar is 20 μm.

Cells induced mitophagy coinciding with mitochondria size reduction. Using Western blot analysis, we observed higher molecular weight bands for an internally 3HA-tagged variant of yeast mitophagy receptor Atg32 after 4 h of starvation, indicating Atg32 phosphorylation and activation of receptor-mediated mitophagy (**Fig. 1A**)(*7, 9, 10*). Consistently, we started to detect mitochondria turnover after 4 h of starvation by measuring generation of free GFP after vacuolar transfer by autophagy and cleavage of mitochondria-bound Om45-GFP (**Fig. 1A**)(*7*). To test whether mitophagy determines mitochondria size dynamics during starvation, we monitored mitochondrial volume in cells deficient for Atg32 (Δ*atg32*). Mitochondrial networks in WT and Δ*atg32* cells were indistinguishable during growth and similarly increased in volume during the first 6 h of starvation, indicating initial mitochondria network expansion is predominantly driven by biogenesis independent of mitophagy (**Fig. 1A**). Strikingly, mitochondria in Δ*atg32* cells continued to expand after 6 h until they reached a roughly two-fold larger volume at 20-24 h starvation compared with WT cells (**Fig. 1A**). These data demonstrate an essential function for Atg32-mediated mitophagy in controlling mitochondrial network size during starvation. Interestingly, in both, WT and Δ*atg32* cells, mitochondrial volume dynamics were determined by changes in mitochondrial tubule length over time while maintaining constant average width (**Fig. S1A**). Notably, the number of nodes (branching points) and tips changed proportionately to total network length, indicating a constant relation between these parameters for mitochondria in WT and Δ*atg32* cells over time (**Fig. S1A**). Thus, mitophagy critically controls total network size with some structural features of mitochondria being unaffected by mitophagy.

To analyze the relationship between autophagy and mitochondria size, we compared the mitochondrial volume of WT cells to cells deficient in all forms of autophagy (Δ*atg7*) and to cells lacking either the adaptor Atg11 for selective forms of autophagy (Δ*atg11*) or the mitophagy receptor Atg32 (Δ*atg32*) (*7, 9, 11*). In line with undetectable levels of autophagy/mitophagy in WT cells monitored by Om45-GFP turnover, neither of the autophagy gene deletions affected mitochondria size during growth (0 h)(**Fig. 1B**). Strikingly, consistent with the established functions of Atg11 and Atg32 in mitophagy and strong defects in mitophagic Om45-GFP turnover, Δ*atg11* and Δ*atg32* cells showed similar increases in mitochondrial volume driven by tubular elongation after 24 and 48 h of starvation (**Fig. 1B** **and S1B**). Notably, Δ*atg7* cells, deficient in both bulk and selective autophagy, did not show a change in mitochondrial volume despite complete absence of detectable Om45-GFP turnover (**Fig. 1B**). Instead, consistent with prior work (*12*), mitochondria quantitatively fragmented during starvation in Δ*atg7* cells illustrated by significantly higher numbers of tips/nodes in total and per unit of mitochondrial length compared with WT cells (**Fig. 1B** **and S1B**). These data uncover that Atg11-Atg32-mediated mitophagy controls mitochondrial network size, and mitochondrial biogenesis and homeostasis depends on bulk autophagy during starvation.

We tested whether mitophagy is not only required but also sufficient to reduce mitochondria size after expansion during starvation. We engineered Δ*atg32* cells to express plasmid-encoded Atg11 fused to FRB (*ATG11^FRB^*) and a mitophagy-inactive Atg32 variant carrying a N-terminal Frp1 domain (*FRP1-atg32^S114,119A^*, designated as *atg32^FRP1^*)(**Fig. 1C**)(*10*). Expressed in a rapamycin-insensitive TORC1 background (*tor1-1*Δ*fpr1*)(*13*), these variants are physically tethered to each other in the presence of rapamycin, allowing for specific and inducible activation of mitophagy (**Fig. 1C**). First, we starved WT, Δ*atg32*, and Δ*atg32* cells harboring an empty vector (*EV*) or the *ATG11^FRB^*-*atg32^FRP1^* construct for 24 h. Except for WT cells, all strains displayed a two-fold increase in mitochondrial volume (**Fig. 1C**). Then we starved cells for an additional 24 h (48 h) in the presence or absence of rapamycin. Untreated *EV* or *ATG11^FRB^-atg32^FRP1^* cells and rapamycin treated *EV* cells maintained expanded mitochondria after 48 h (**Fig. 1C**). Strikingly, rapamycin-treatment specifically induced mitophagy in *ATG11^FRB^-atg32^FRP1^* cells, which reduced expanded networks to the size of WT mitochondria (**Fig. 1C**). Taken together, these data demonstrate that Atg11-Atg32-mediated mitophagy is required and sufficient for pruning expanded mitochondrial networks to pre-starvation size.

To test whether the role of receptor-mediated mitophagy in mitochondrial network size regulation is evolutionarily conserved, we efficiently knocked down the mammalian homolog of Atg32, Bcl2L13, in mouse embryonic fibroblasts (MEFs) by siRNA treatment (**Fig. S1C**)(*14*). MEFs were grown in full media and mitochondrial networks were visualized by Mitotracker staining and fluorescence microscopy followed by MitoGraph analysis (*15*). Strikingly, compared with untreated or scrambled siRNA controls, mitochondrial volume significantly increased upon Bcl2L13 knock down (**Fig. 1D**). These data are consistent with an evolutionarily conserved function of Atg32/Bcl2L13-mediated mitophagy in controlling mitochondrial network size from yeast to mammals.

### Mitochondria maintain wildtype-like features in the absence of mitophagy

Mitophagy has been linked to quality control of mitochondria by selectively degrading dysfunctional portions of the network (*4, 16*). We asked whether mitochondrial expansion in the absence of mitophagy reflected impaired quality control and as a consequence resulted in the accumulation of dysfunctional mitochondria. Alternatively, and not mutually exclusively, defects in mitophagy or mitophagy receptors may cause an altered cellular metabolic state and trigger mitochondria biogenesis. To address these questions, we profiled global proteome changes in dependence of autophagy (Δ*atg7*), selective autophagy (Δ*atg11*), or mitophagy (Δ*atg32*) using multiplexed tandem mass tag (TMT) whole cell proteomics. Defects in autophagy caused broad starvation-induced differences in the global and mitochondrial proteome of Δ*atg7* cells compared with WT cells (**Fig. 2A and B**), indicating significant autophagy-dependent proteome remodeling upon nutrient stress. In contrast, global proteomes were much less impacted in Δ*atg11* and Δ*atg32* cells than in Δ*atg7* cells. Notably, we observed a ∼1.3-fold average increase of mitochondrial proteins in Δ*atg11* and Δ*atg32* cells relative to WT cells after starvation (**Fig. 2A**), consistent with increased mitochondrial volume and protein mass in the absence of mitophagy. When normalized to average protein abundance, the relative mitochondrial proteome composition of Δ*atg11* and Δ*atg32* cells strongly correlated with WT cells in contrast to Δ*atg7* cells (**Fig. 2B**). These data show that defects in mitophagy cause elevated mitochondrial protein content in cells alongside larger mitochondria volume. However, mitophagy-deficient cells maintain a largely wildtype-like mitochondrial proteome composition.

**Figure 2:**
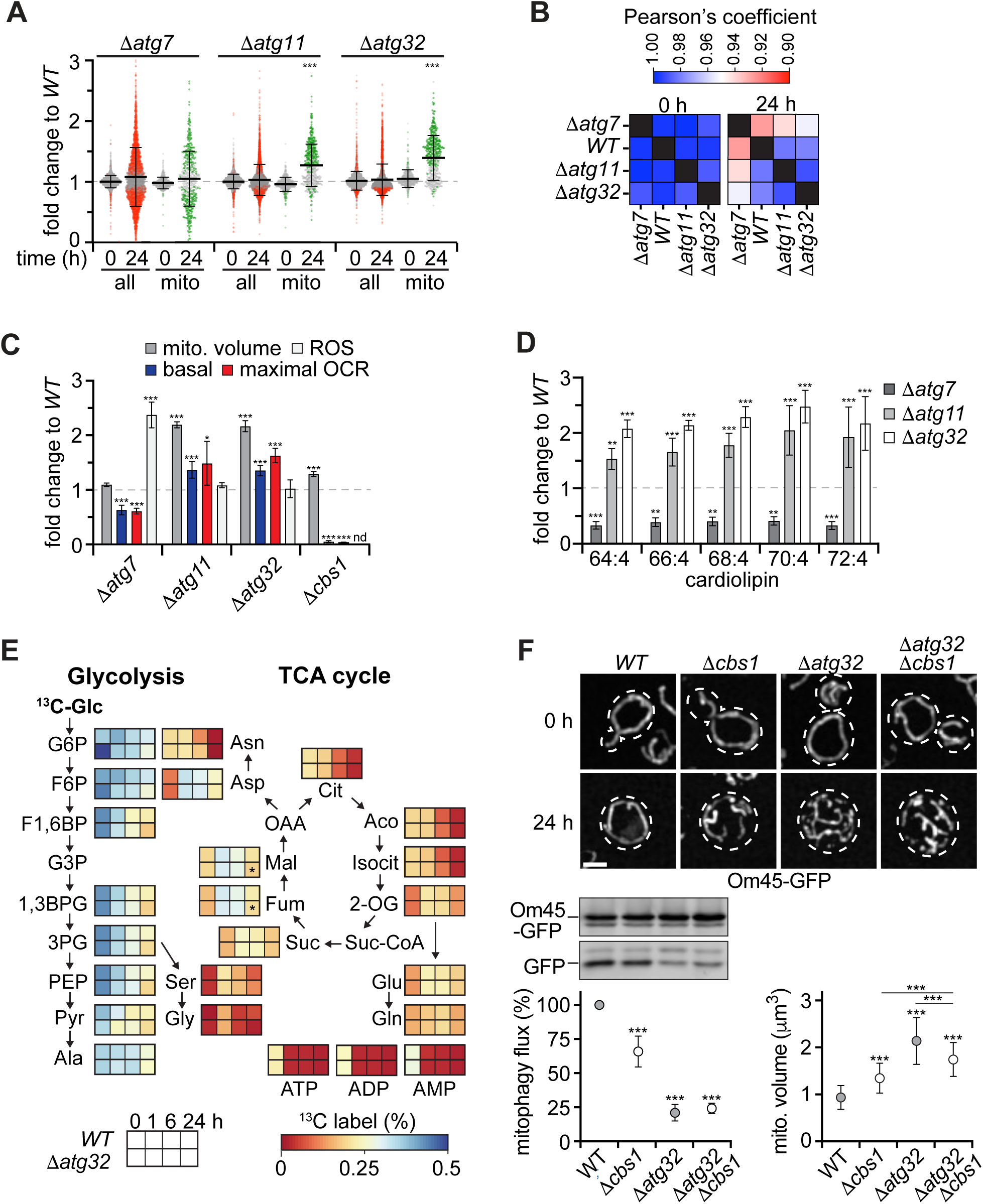
Mitochondria maintain wildtype-like features in the absence of starvation-induced mitophagy. **(A)** Whole cell multiplexed tandem mass tag (TMT) proteomics of indicated strains before (0 h) and after starvation (24 h). 3759 proteins in total and 565 mitochondrial proteins were quantitatively analyzed. Proteins significantly changed in indicated cells compared with WT cells are marked in red or green for all or mitochondrial proteins, respectively. Means ± SD are shown. **(B)** Similarity of normalized mitochondrial protein composition for indicated strains before (0 h) and after starvation (24 h). **(C)** Mitochondrial function based on volume, basal and maximal oxygen consumption rate (OCR) or production of reactive oxygen species (ROS) in indicated strains after starvation (24 h). **(D)** Lipidomics-based cardiolipin levels of whole indicated cells. **(E)** ^13^C-glucose tracing in WT and Δ*atg32* cells at indicated time points during starvation. ^13^C-incorporation rates (^13^C/^12^C in %) for indicated metabolites are shown. Data are means ± SD (n=5). **(F)** Mitochondrial volumes and mitophagy for indicated strains expressing *OM45*-GFP before (0 h) and after starvation (24 h). Fluorescence imaging (upper panel), Western blot analysis of Om45-GFP turnover in cell lysates using α-GFP antibody (lower left panel), and Mitograph analysis of mitochondria volume (lower right panel). Representative average intensity Z-stack projections are shown. Data are means ± SD (mitophagy flux: n=6; mitochondria volume: n=3, 60 cells/data point). Scale bar is 2 μm.

We assessed the functional state of mitochondria in dependence of autophagy and mitophagy. We measured production of reactive oxygen species (ROS) and basal or maximal oxygen consumption rates (OCR). Autophagy-defects caused a strong increase in ROS production and a significant decrease in basal and maximal OCR in Δ*atg7* cells compared with WT cells (**Fig. 2C**), indicating that cells develop mitochondrial dysfunction during starvation in the absence of autophagy. In contrast, Δ*atg11* and Δ*atg32* cells produced WT-like ROS levels despite a ∼2-fold increase in mitochondrial volume (**Fig. 2C**). Δ*atg11* and Δ*atg32* cells consumed oxygen at higher basal and maximal rates compared with WT cells, but not to scale to increased mitochondrial volume (**Fig. 2C**), indicating reduced respiration and ROS production per unit of mitochondrial volume. Interestingly, elevated respiration correlated with increased mitochondrial protein content (∼1.3-fold each) in Δ*atg11* and Δ*atg32* cells (**Fig. 2A and C**). These data suggest similar respiration to mitochondrial protein ratios in WT and mitophagy-deficient cells and are fully consistent with the observed increase in mitochondrial protein content without significant compositional changes in the absence of mitophagy (**Fig. 2A-C**). Because the mitochondrial proteome did not increase to scale with mitochondrial volume in mitophagy-deficient cells, we performed whole cell lipidomics using mass spectrometry to measure the content of cardiolipin, a signature lipid of mitochondria, as a proxy for mitochondrial membrane content. Interestingly, the most abundant cardiolipin forms strongly decreased in Δ*atg7* cells after starvation (24 h)(**Fig. 2D**), paralleling starvation-induced morphological and functional decline of mitochondria in the absence of autophagy. In contrast, Δ*atg11* and Δ*atg32* cells showed a 1.5- to 2-fold increase in cardiolipins compared with WT cells (**Fig. 2D**), suggesting proportionate increase of mitochondrial lipid content with organelle volume (**Fig. 2C and D**). To test whether expanded mitochondrial networks in Δ*atg32* cells might cause or are a consequence of metabolic alterations, we traced ^13^C-glucose metabolism using mass spectrometry (**Fig. 2E**). Notably, WT and Δ*atg32* cells displayed virtually indistinguishable metabolic rates and changes in glycolysis and tricarboxylic acid (TCA) cycle during growth (0 h) and at different time points of starvation (1, 6, or 24 h)(**Fig. 2E**), indicating that expanded networks are neither a result of nor cause significant metabolic changes in mitophagy-deficient cells. Taken together, our data show that expanded mitochondrial networks in mitophagy-deficient cells maintain wildtype-like structural, compositional, and functional features.

Our data indicate that, in contrast to defects in autophagy, mitophagy-deficient cells do not accumulate detectable levels of mitochondrial dysfunction. This suggests that starvation-induced mitophagy does not function primarily in mitochondrial quality control. To critically challenge this notion, we examined mitophagy and mitochondrial volume in cells with severe mitochondrial respiratory deficiency. Δ*cbs1* cells lack an essential factor for translation of mitochondrial encoded Cob1, a core subunit of CIII, and are therefore incapable of mitochondrial respiration (**Fig. 2C**)(*17*). Importantly, when starved, Δ*cbs1* cells displayed significantly reduced mitophagy compared with WT cells (**Fig. 2F**). Thus, instead of triggering quality control mitophagy, severe mitochondrial dysfunction suppresses mitophagy during starvation. Consistent with reduced size control by mitophagy, Δ*cbs1* cells displayed a slightly expanded mitochondrial volume compared with WT cells after starvation (**Fig. 2F**). Interestingly, the absence of Cbs1 decreased mitochondrial expansion in mitophagy-deficient cells (Δ*atg32*Δ*cbs1*)(**Fig. 2F**). Taken together, our data suggest that, in response to starvation, mitophagy is homeostatic in nature and functions primarily in mitochondrial size regulation rather than quality control.

### Bulk autophagy sustains mitochondria biogenesis during starvation

We made the unexpected observation that mitochondria undergo significant biogenesis in starving and non-dividing cells, resulting in expanded networks in the absence of mitophagy. Mitochondrial biogenesis depended on autophagy, because the absence of the core autophagy component Atg7 alone (Δ*atg7* cells) or in combination with Atg32 (Δ*atg7*Δ*atg32* cells) significantly suppressed mitochondrial network expansion observed in Δ*atg11* or Δ*atg32* cells (**Fig. 3A**). These data clearly indicate that mitochondrial biogenesis relies on bulk autophagy during starvation.

**Figure 3:**
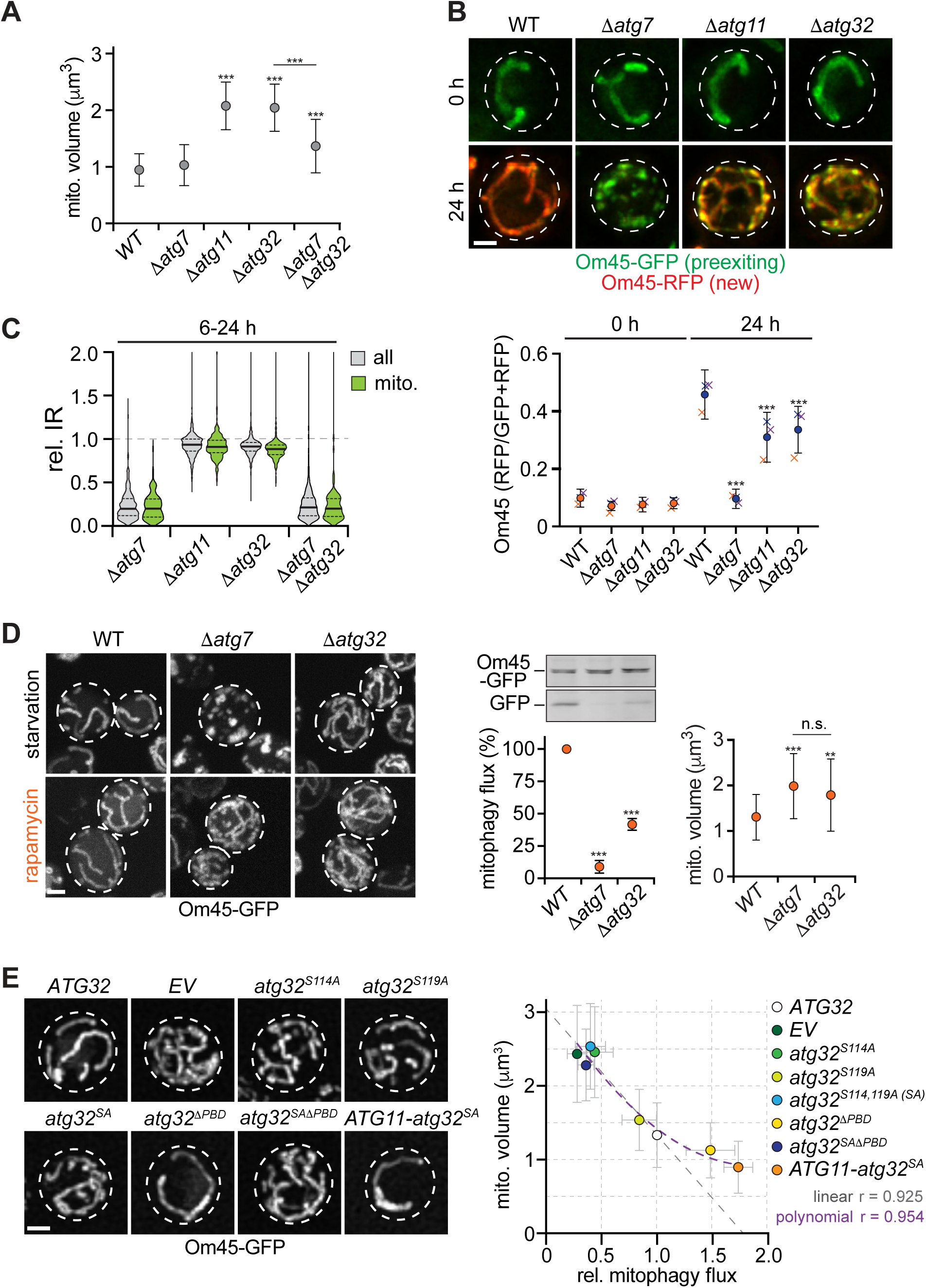
Autophagy sustains mitochondrial biogenesis counterbalancing elevated mitophagy. **(A)** Mitochondria volumes in indicated cells based on fluorescence imaging of MITO-ID Green stained mitochondria and Mitograph analysis. **(B)** Analysis of pre-existing Om45-GFP and newly synthesized Om45-RFP in indicated cells before (0 h) and after starvation (24 h) after recombination induced tag exchange after 6 h of starvation. Representative average intensity Z-stack projections are shown. Data are means ± SD, means for each replica are shown (n=3, 30 cells/replica). **(C)** Global and mitochondrial protein synthesis measured as relative incorporation rate by whole cell tandem mass tag (TMT) proteomics after labeling with isotopic lysine (lys^8^: ^13^C_6_,^15^N_2_) for 6-24 h of starvation. lys^8^/lys^0^ rates were normalized to WT (set as 1) for each identified protein. Data are means (n=4). **(D)** Fluorescence imaging of indicated cells expressing *OM45-GFP* after starvation or rapamycin treatment (24 h). Om45-GFP turnover in indicated cells (middle panel) and mitochondria volumes based on Mitograph analysis (right panel) after rapamycin treatment (24 h). Representative average intensity Z-stack projections are shown. Data are means ± SD (mitophagy flux: n=6; mitochondria volume: n=3, 60 cells/data point). **(E)** Analysis of mitochondria volume and mitophagy flux in Δ*atg32* cells expressing *OM45*-GFP and indicated plasmid encoded variants of *ATG32*. Fluorescence imaging (left panel) and comparison of mitochondria volume and mitophagy flux (right panel). Regressions are based on all strains (polynomial) or except for *atg32*^Δ*PBD*^ and *ATG11-atg32^SA^* expressing strains (linear). Primary data are shown in figure S2C. Representative average intensity Z-stack projections are shown. Data are means ± SD (mitochondria volume: n=3, ≥60 cells/data point; mitophagy flux: n=6). Scale bars are 1 μm.

We tested whether bulk autophagy drives mitochondrial protein synthesis during starvation. We used the recombination induced tag exchange (RITE) system to monitored protein levels of pre-existing Om45-GFP and, after Cre-recombinase induced tag exchange after 6 h of starvation, generation of newly synthesized Om45-RFP during starvation (24 h)(*18*). In WT cells, mitochondria strongly enriched newly synthesized Om45-RFP over pre-existing Om45-GFP during starvation (**Fig. 3B**). Om45-GFP turnover was promoted by mitophagy, because Δ*atg11* and Δ*atg32* cells synthesized Om45-RFP, but maintained higher levels of pre-existing Om45-GFP than WT cells after 24 h starvation (**Fig. 3B**). Strikingly, Δ*atg7* cells failed to express detectable levels of Om45-RFP and mitochondria remained almost exclusively marked by Om45-GFP (**Fig. 3B**), demonstrating cells require bulk autophagy for Om45 translation during starvation. Whole cell proteomics after isotopic labeling (SILAC) confirmed protein synthesis depends on bulk autophagy, because Δ*atg7* and Δ*atg7*Δ*atg32* cells showed drastically decreased incorporation of isotopic lysine (lys^8^) into newly synthesized proteins at global and mitochondrial protein level during starvation compared with WT, Δ*atg11*, or Δ*atg32* cells consistent with prior work (**Fig. 3C**)(*19*). Interrogating the substrate scope of bulk autophagy, we observed a significant 2-fold decrease in the steady-state levels of ribosomal proteins in WT cells after starvation (24 h) in dependence of bulk autophagy in Δ*atg7* cells, but not of selective autophagy or mitophagy in Δ*atg11* or Δ*atg32* cells, respectively (**Fig. S2A**). These data indicate that yeast cells degrade about half of their ∼200.000 ribosomes in a bulk autophagy-dependent manner within the first 24 h of starvation (*20*). Interestingly, cells maintained steady-state levels of proteasomal proteins after starvation independent of bulk or selective autophagy (**Fig. S2A**), suggesting differential effects of starvation-induced autophagy on large cytosolic complexes. This apparent selectivity for ribosome degradation likely arises from bulk autophagy of both, ribosomes and proteasomes, combined with differential translation of proteasomal but not ribosomal proteins during starvation (**Fig. S2A and B**). In combination, these data indicate that bulk autophagy sustains translational programs of starving cells with considerable ribosome degradation underlying mitochondrial biogenesis. In strong support of this conclusion, induction of autophagy by rapamycin-treatment in the presence of nutrients caused mitochondrial expansion in autophagy-deficient Δ*atg7* cells indistinguishable from Δ*atg32* cells due to a complete block in mitophagy but sustained mitochondrial biogenesis in contrast to starvation (**Fig. 3D**). Thus, nutrient availability renders bulk autophagy dispensable for mitochondrial biogenesis, but mitophagy remains essential for size control of mitochondria networks. Taken together, these data indicate that mitophagy acts downstream of bulk autophagy-driven mitochondrial biogenesis. Bulk autophagy liberates metabolites critical for mitochondrial biogenesis during starvation, and resulting mitochondrial expansion triggers mitophagy to control mitochondrial network size.

### Mitochondrial biogenesis compensates for synthetically elevated mitophagy

Our data indicate that cells induce mitophagy to prune mitochondrial networks in response to biogenesis-driven expansion. We asked whether cells may also sense and respond to decreased mitochondrial volume due to elevated mitophagy. To address this question, we measured mitophagy flux by Om45-GFP turnover in correlation with mitochondrial volume changes in Δ*atg32* cells expressing a set of plasmid-encoded Atg32 variants promoting different levels of mitophagy (**Fig. 3E** **and S2C**). Strong reduction in mitophagy correlated with a ∼2-fold volume increase of mitochondria in cells expressing none (*EV*) or inactive *atg32^S114A^*or *atg32^S114,119A^* variants, which are defective in phosphorylation-regulated Atg11 binding, compared with wildtype *ATG32* (**Fig. 3E** **and S2C**)(*10*). Interestingly, the presence of the slightly less active *atg32^S119A^* variant caused a mild but proportionate increase in mitochondrial volume compared with wildtype or inactive *ATG32* variants (**Fig. 3E** **and S2C**). These data predict a linear relation between mitochondrial volume and the level of mitophagy modeled by linear regression (**Fig. 3E**). To synthetically elevate mitophagy, we expressed an Atg32 variant deficient for the binding of the inhibitory phosphatase Ppg1, resulting in constitutive phosphorylation and activation of Atg32^ΔPBD^ in contrast to a non-phosphorylatable variant carrying S114A and S119A mutations (Atg32^SAΔPBD^)(*10, 21*). In addition, we designed a hybrid protein of Atg11 fused to inactive Atg32^S114,119A^ (*ATG11-atg32^SA^*), constitutively tethering Atg11 to Atg32 on the outer membrane of mitochondria. Expression of *atg32*^Δ*PBD*^ or *ATG11-atg32^SA^* resulted in significantly 1.5-or 1.7-fold increased mitophagy compared to *ATG32* cells, respectively (**Fig. 3E** **and S2C**). Interestingly, these elevated levels of mitophagy did not decrease mitochondrial volumes according to linear extrapolation (**Fig. 3E** **and S2C**). Instead, these cells displayed mitochondrial volumes approaching an apparent minimal mitochondrial volume described by polynomial regression (**Fig. 3E**). These data suggest that cells counteract increased mitophagy by compensatory biogenesis to maintain minimal mitochondria volume. Thus, we propose that cells respond to changes in mitochondria size not only by inducing mitophagy due to increased volume but also by upregulating mitochondrial biogenesis in the presence of excess mitophagy.

### Metabolically tunable setpoints define mitochondria scaling by autophagy

Our work establishes a dual role for autophagy in size regulation of mitochondria during starvation with bulk autophagy driving biogenesis and homeostatic mitophagy reducing expanded networks to pre-starvation size. However, cellular parameters or “setpoints” to which starving cells scale mitochondria volume to are unknown. To begin to identify potential setpoints for mitochondria scaling by autophagy, we analyzed in parallel the size parameters of cell, vacuole, cytosol, and mitochondria volume at single-cell resolution using fluorescence imaging of cells with Om45-GFP-marked mitochondria and vital dye FM4-64-stained vacuole membranes. We observed a significant increase in cell volume in WT, Δ*atg7*, Δ*atg11*, and Δ*atg32* cells after starvation (24 h) (**Fig. S3A**). Starvation-induced increase in cell volume was concomitant with an enlargement of vacuoles (**Fig. S3B)**. Interestingly, Δ*atg7* cells displayed a smaller cell and vacuole volume before and after starvation compared with WT cells (**Fig. S3A**). However, WT, Δ*atg11*, and Δ*atg32* cells showed similar vacuole volume ratio changes and isometric relation of the vacuole-to-cell volume at single-cell level upon starvation consistent with prior work (**Fig. S3E and S4**)(*22*). Interestingly, cytosol volume, approximated as non-vacuolar cell volume, remained the same before and after starvation in all tested strains (**Fig. S3D**). Taken together, these data indicate that cells, while maintaining constant cytosol volume, increased in total volume due to vacuolar expansion in response to starvation.

Previous work connected mitochondrial network size to an increasing cell volume based on a strong correlation or scaling relation in growing buds of dividing yeast cells in the presence of nutrients (*6*). When we interrogated the relation between mitochondria and cell volume in starving cells, we observed only a moderate correlation at single-cell level (**Fig. S4**). According to prior definition (*23*), we considered scaling setpoints as mean values of size parameters at population level. Interestingly, in the absence of mitophagy (Δ*atg11* and Δ*atg32* cells), we observed mitochondria volume increase proportionate with enlarged cell volume (**Fig. 4A**). Consistently, analysis of mitochondria and cell volume for the time course of 24 h, as shown in figure 1A, uncovered a strong correlation between the average mitochondria and cell volume and a constant mitochondria volume ratio after 2 h of starvation in Δ*atg32* cells (**Fig. S5A and B**). These data indicate that, driven by mitochondria biogenesis, mitochondria volume scales with cell volume in the absence of mitophagy, suggesting the existence of a mitochondria-to-cell volume setpoint. Consistent with a phase of predominant biogenesis, mitochondria volume also scaled with cell volume in WT cells with a mitochondria volume ratio similar to Δ*atg32* cells during the first 6-8 h of starvation (**Fig. S5A and B**). Strikingly, upon mitophagy-dependent reduction of mitochondria volume after 8 h, WT cells displayed a significantly different mitochondria-to-cell volume ratio at 24 h of starvation compared with WT cells before starvation (0 h) and with mitophagy-deficient Δ*atg11* and Δ*atg32* cells (**Fig. 4A, S5A and B**). These data suggest the existence of additional size setpoint(s) for scaling of WT mitochondria by mitophagy during starvation. Notably, although only moderately correlated at single-cell level (**Fig. S4**), we observed that WT cells reestablished the same mitochondria to cytosol volume ratio at population level after starvation (24 h) as during growth (0 h) (**Fig. 4A** **and S3G**). These data suggest that WT cells use mitophagy to scale mitochondria volume according to a mitochondria-to-cytosol volume setpoint. Taken together, our data suggest a model in which mitochondria biogenesis initially increases mitochondria volume according to a mitochondria-to-cell volume setpoint. Then, mitophagy is triggered to scale mitochondria volume according to a mitochondria-to-cytosol volume setpoint.

**Figure 4:**
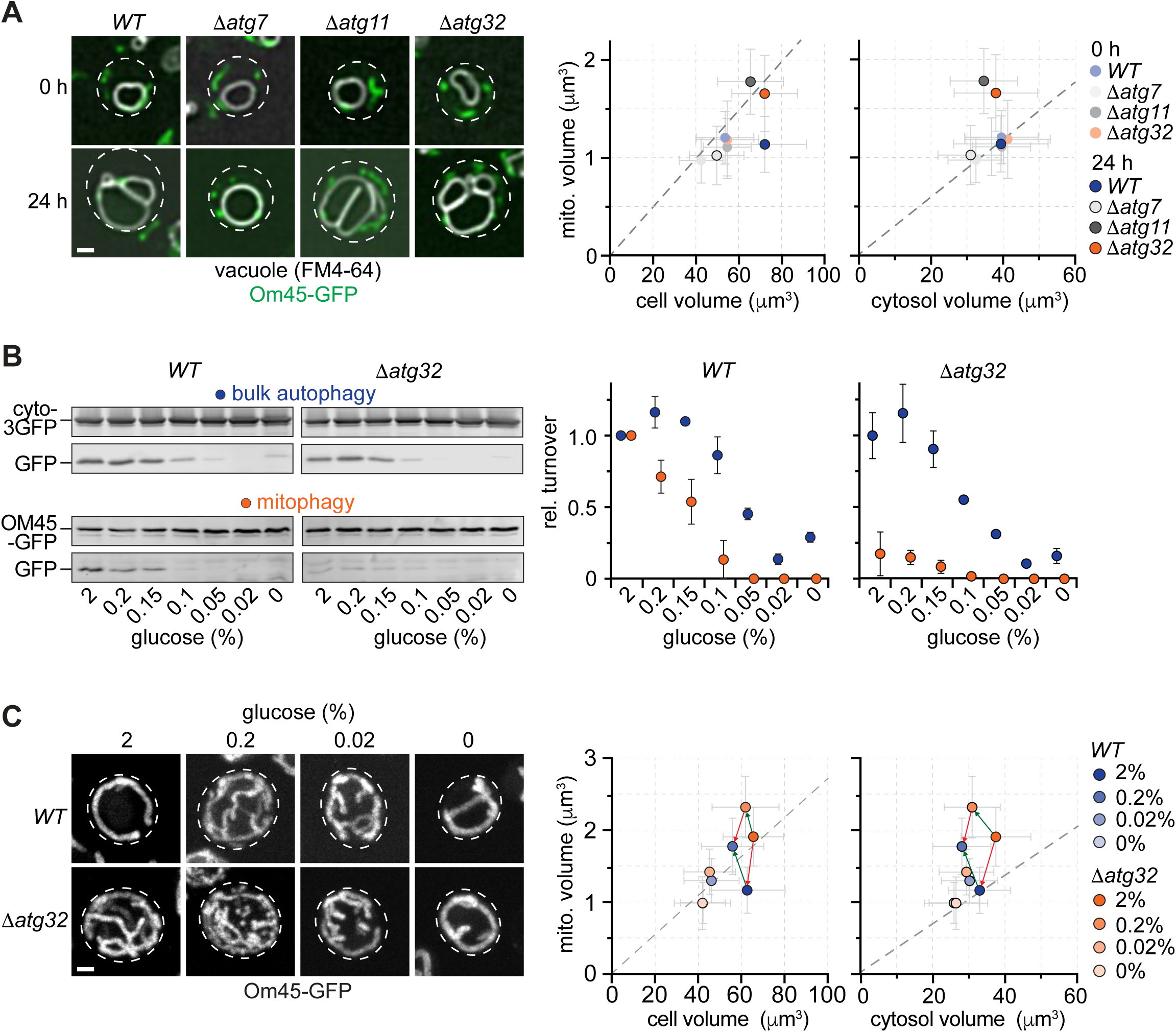
Single-cell analysis identifies metabolically tunable setpoints for mitochondria scaling by autophagy. **(A)** Analysis of cell, vacuole, cytosol, and mitochondria volume at single-cell level by fluorescence imaging in indicated cells before (0 h) and after starvation (24 h). *OM45-GFP* expressing cells were treated with FM4-64 to visualize vacuolar membranes. Fluorescence imaging (left panel) and comparison of mitochondria volume to cell or cytosol volume (middle and right panel, respectively) are shown. Representative average intensity Z-stack projections are shown. Data are means ± SD (n=3, ≥145 cells/data point). **(B)** Analysis of cytosolic 3GFP (bulk autophagy) and Om45-GFP (mitophagy) turnover in indicated cells in dependence of glucose availability. **(C)** Analysis of mitochondria to cell or cytosol volume in dependence of glucose availability in WT and Δ*atg32* cells expressing *OM45*-GFP and stained with FM4-64 after starvation (24 h). Fluorescence imaging (left panel) and comparison of mitochondria and cell or cytosol volume (middle and right panel, respectively) are shown. Representative average intensity Z-stack projections are shown. Data are means ± SD (n=3, ≥150 cells/data point). Scale bars are 1 μm.

We asked whether cells use mitophagic scaling to adapt mitochondria volume to metabolic inputs. To address this question, we exposed cells to decreasing concentrations of glucose during starvation. Glucose is a central metabolite, which suppresses (2%), de-represses (0.2%) or induces mitochondrial biogenesis in growing cell populations. We established a cytosolic 3GFP variant as a reporter for bulk autophagy (**Fig. S5C**). Next, we measured the effects of different glucose concentrations on cytosolic (cyto-3GFP) and mitochondrial (Om45-GFP) turnover in WT and Δ*atg32* cells. Interestingly, decreasing glucose concentrations correlated with reduced cytosol turnover in WT and Δ*atg32* cells (**Fig. 4B**), indicating that glucose is limiting for bulk autophagy induction in starving cells. Similarly, the levels of mitophagy strongly declined in WT cells exposed to lower glucose concentrations (**Fig. 4B**), showing that both, bulk and mitophagy depend on glucose availability during starvation. Next, we monitored the changes in cell, vacuole, cytosol, and mitochondria volume in WT and Δ*atg32* cells with Om45-GFP-marked mitochondria and vital dye FM4-64-stained vacuole membranes exposed to different concentrations of glucose during starvation (**Fig. S6**). Strikingly, in the presence of 0.2% glucose, WT cells displayed a significantly increased mitochondria volume in line with slightly increased bulk autophagy and reduced mitophagy compared with WT cells exposed to 2% glucose (**Fig. 4C** **and S6**), indicating that cells alter mitophagic scaling to adjust the mitochondria volume to a different mitochondria-to-cytosol volume setpoint. In addition, Δ*atg32* cells also developed an increased mitochondria volume (**Fig. 4C**), indicating that cells adapt a new mitochondria-to-cell volume setpoint. Thus, although cells increase mitochondrial biogenesis in the presence of 0.2 vs 2% glucose during starvation as seen for Δ*atg32* cells, they still use mitophagic scaling to adjust mitochondria volume. Interestingly, when exposed to 0.02 or 0% glucose, we did not observe a detectable difference in mitochondria volume between WT and Δ*atg32* cells (**Fig. 4C**). Consistently, we did not detect any differences in bulk autophagy or, moreover, any mitophagy in WT and Δ*atg32* cells (**Fig. 4B**). These data suggest that 0.02% glucose sustains limited mitochondrial biogenesis in both WT and Δ*atg32* cells compared with 0% glucose, but the increase in mitochondria volume is insufficient to induce mitophagy. Taken together, these data indicate that cells regulate autophagy responses composed of bulk autophagy and mitophagy to scale mitochondria volume according to two metabolically tunable setpoints, mitochondria-to-cell and -cytosol volume, to integrate cellular size and metabolic parameters.

Our work identifies a role for homeostatic mitophagy in scaling mitochondria network size in yeast and mammals. Understanding the molecular interplay of quality control mitophagy and mitophagic scaling may pave the way for novel therapeutic strategies for mitochondrial diseases.

## Acknowledgments

The authors would like to the thank the Imaging and FACS, Metabolomics, and Proteomics core facilities of the MPI-AGE for outstanding support.

## Funding

This work was supported by the Max Planck Society (MPG) and the Deutsche Forschungsgemeinschaft (DFG-German Research Foundation) – SFB 1218 – project number 269925409 to M. Graef.

## Author contributions

Conceptualization: RLT, MG

Investigation and Validation: RLT, BCB, LA, MG

Formal analysis: HN

Supervision: MG

Funding acquisition: MG

Visualization: RLT, BCB, LA, NH, MG

Writing original draft: MG

Writing – editing and review: RLT, BCB, LA, HN, MG

## Competing interests

Authors declare that they have no competing interests.

---

**Table S1:**
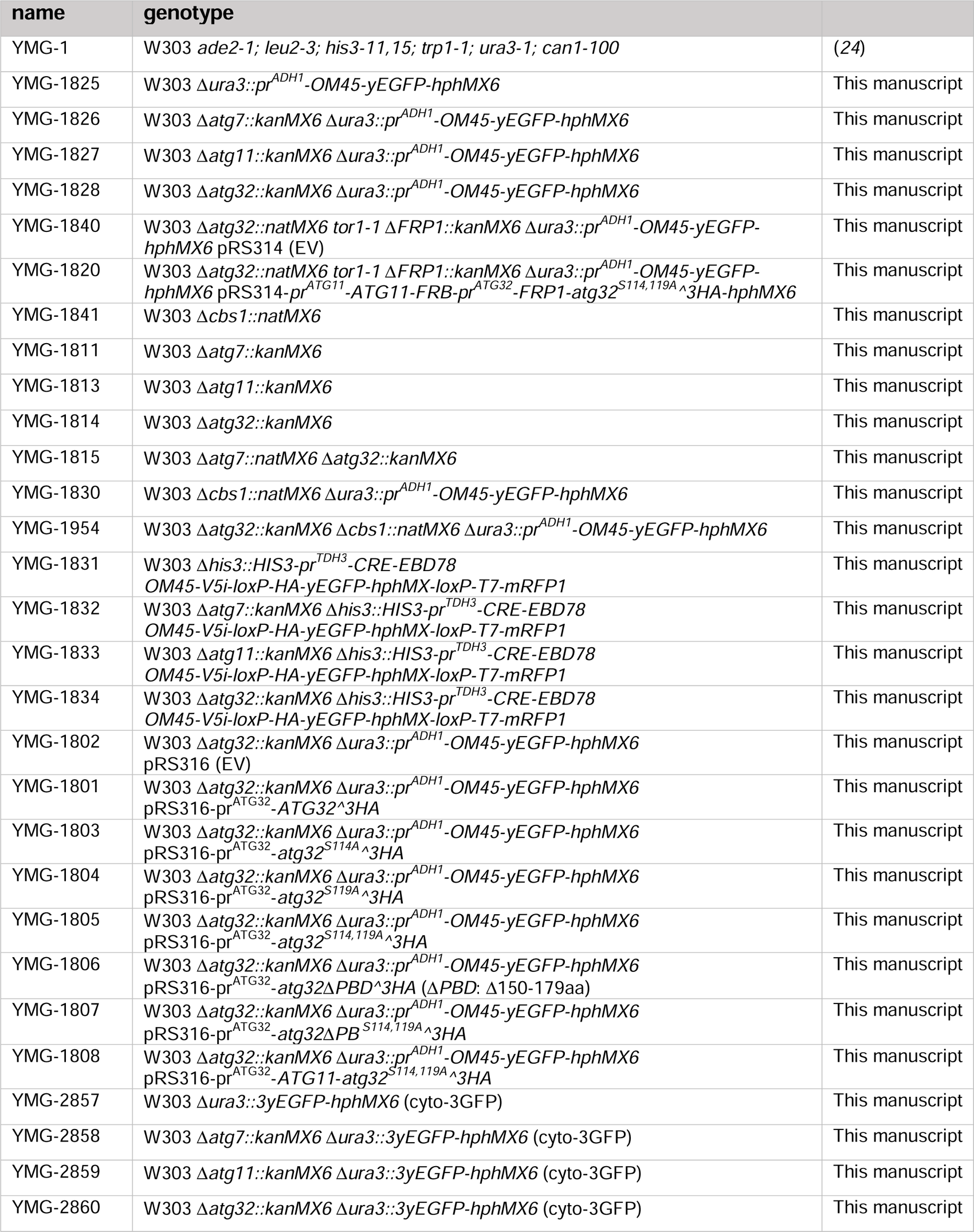
Yeast strains used in this study.

## Materials and Methods

### Yeast strains

Yeast strains used in this study are W303 or derivatives thereof and are listed in **Table S1**. Gene deletions were generated by replacing complete open reading frames (ORFs) with indicated PCR-amplified marker cassettes using targeted homologous recombination as described previously (*25*). For the *OM45-GFP* reporter, the ADH1 promoter (pr^ADH1^: 700 bp 5’ upstream of *ADH1*) was fused to *OM45* without stop codon linked to *linker-yEGFP* (amplified from pFA6a-yEGFP-CaURA3) and the *hphMX6* cassette (pFA6a-*hphMX6*). The construct was integrated into the genome replacing the *URA3* locus using targeted homologous recombination. For the cyto-3GFP reporter, *3yEGFP* was amplified from pFA6a-*3yEGFP-CaURA3* (*26*) and integrated into the genome replacing the *URA3* locus using targeted homologous recombination fused to the *hphMX6* cassette (pFA6a-*hphMX6*). The plasmid pRS316-*pr^ATG32^-ATG32∧3HA* series was derived from pRS315*-pr^ATG32^-ATG32∧3HA* described previously (*9*) and the following changes were introduced individually or in combination: S114A (TCA to GCC), S119A (TCT to GCC), and ΔPBD (deletion of aa 150-179). The plasmid pRS316-*pr^ATG32^-ATG11-atg32^S114,119A^∧3HA-hphMX6* was generated by gap repair cloning fusing *ATG32* promoter (pr^ATG32^: 580 bp 5’ upstream of *ATG32*), *ATG11* without stop codon, *atg32^S114,119A^∧3HA*, ATG32 3’UTR (193 bp 3’ downstream of *ATG32*) and *hphMX6* carrying the *PGK1* promoter and terminator (pr^PGK1^: 1 kb 5’ upstream of *PGK1*; ter^PGK1^: 271 bp 3’ downstream of *PGK1*). The plasmid pRS314-*pr^ATG11^-ATG11-FRB-pr^ATG32^-FRP1-atg32^S114,119A^∧3HA-hphMX6* was generated by gap repair cloning fusing *ATG11* promoter (pr^ATG11^: 279 bp 5’ upstream of ATG11), *ATG11* without stop codon, *FRB*, ATG11 terminator (ter^ATG11^: 300 bp 3’ downstream of *ATG11*), *ATG32* promoter (pr^ATG32^: 580 bp 5’ upstream of *ATG32*), *FRP1* without stop codon, *atg32^S114,119A^∧3HA*, ATG32 3’UTR (193 bp 3’ downstream of *ATG32*) and *hphMX6* carrying the *PGK1* promoter and terminator (pr^PGK1^: 1 kb 5’ upstream of *PGK1*; ter^PGK1^: 271 bp 3’ downstream of *PGK1*).

### Media

Yeast cells were grown in synthetic defined complete (SDC) containing 2% (w/v) α-D-glucose (Sigma), 0.7% (w/v) yeast nitrogen base (BD Difco) and a complete supplement of amino acids and nucleobases. Strains carrying indicated plasmids were grown in the absence of the respective metabolite corresponding to auxotrophic markers. For nitrogen starvation, cells were grown in SDC overnight to early log-phase and then washed at least three times in synthetic defined without nitrogen (SD-N) media composed of 0.17% (w/v) yeast nitrogen base without amino acids and ammonium sulfate (BD Difco) and 2% (w/v), as standard, or 0.2, 0.15, 0.1, 0.05, 0.02 or 0% (w/v) indicated concentrations of α-D-glucose (Sigma). To maintain constant osmolarity, SD-N media containing >2% (w/v) α-D-glucose were supplemented with corresponding concentrations of D-sorbitol to a maintain total concentration of 2% (w/v)(Figure 4B and C). For induction of the FRB-FRP1 tether (Figure 1C) or of autophagy in SDC media (Figure 3D), a final concentration of 50 μM rapamycin (50 mM stock in DMSO) was added to the media. Equal volumes of DMSO were added to control samples.

### Mammalian cell culture, siRNA knockdown and fluorescence microscopy

Mouse Embryo Fibroblast cells (MEFs, NIH-3T3) were cultured in Dulbecco’s Modified Eagle Medium (DMEM) supplemented with 10 % (v/v) Fetal Bovine Serum, 1 % (v/v) penicillin-streptomycin (10.000 U/ml), 1 % (v/v) GlutaMAX and 1 % (w/v) sodium pyruvate at 37 °C and 5% CO_2_. Wt MEF cells were transfected with the respective ON-TARGETplus siRNAs (Horizon; SCR control: D-001810-10; Bcl2L13: J-054573-08) using Lipofectamine RNAiMAX (Invitrogen) according to the manufacturer’s protocol. Briefly, 24 h before transfections, 10,000 cells were seeded in 24-well plates to 60% confluency. An siRNA-lipid complex, comprising 25 nmol of the siRNA, in 25 μL of Opti-MEM mixed with 1.5 μL of Lipofectamine RNAiMAX in 25 μL of Opti-MEM was prepared. After incubation for 5 min at room temperature, cells were incubated for 24 h with the siRNA-lipid complex and then left in complete medium for another 48 h before analysis.

Knockdown was validated by qPCR analysis: RNA was isolated using the Aurum Total RNA Mini Kit (Bio-Rad). RNA yield and purity were verified by absorbance at 260/280 nm (NanoVUE spectrophotometer, GE Healthcare Life Sciences). 500 ng RNA was reverse transcribed using the iScript cDNA Synthesis Kit (Bio-Rad) according to the manufacturer’s instructions. For quantitative RT-PCR (RT-qPCR), 3 μl ten-fold diluted cDNA preparation was mixed in a final volume of 10 μl containing 5 μl iQ SYBR Green Supermix (Bio-Rad) and 4 pmol of each primer (Fw: 5’-AGTGGAGACTGCAGTCCATG-3’; Rw: 5’-ACTCTGCGGCGTGCTC-3’). Samples were analysed using a CFX384 Real-time System (Bio-Rad). A melting curve was obtained for each sample in order to confirm single product amplification. cDNA samples from no template control (NTC) and no reverse transcriptase control (NRT) were included as negative controls. GAPDH (Fw: 5’-GTCGGTGTGAACGGATTTG-3’; Rw: 5’-GAACATGTAGACCATGTAGT

### TG-3’) and HPRT1 (Fw: 5’-CCACAGGACTAGAACACCTGCTAA-3’; Rw: 5’-GCCCTGAGG

CTCTTTTCCAG-3’) were used as reference genes. MEFs were grown on coverslips (VWR; 12 mm diameter) and then fixed with 4% (v/v) paraformaldehyde (VWR) at room temperature for 1h. Fixed cells were washed three times with 1×PBS for 10 min and then further permeabilized by incubation with 1% (v/v) Triton X-100 and 1% (w/v) glycin for 10 min. Cells were washed three times with 1×PBS, blocked with 5% (w/v) bovine serum albumin (BSA) solution at room temperature for 1 h, and stained with DAPI (1:10,000) for 10 min at room temperature. Cells were washed with 1×PBS three times. Coverslips were mounted on glass slides with ProLong Gold (ThermoFisher) and imaged using widefield fluorescence imaging was on a Leica DMI6000B with LASX software. Post-processing of the images was done using ImageJ software (v. 3.10).

### Live cell fluorescence imaging

Cells were transferred to 96-well glass bottom microplates (Greiner Bio-One) from indicated cultures and allowed to settle for 15 min. Cells were imaged at room temperature with an inverted microscope (Nikon Ti-E) using a Plan Apochromat IR 60x 1.27 NA objective (Nikon) and Spectra X LED light source (Lumencor). Three-dimensional light microscopy data were collected using the triggered Z-Piezo system (Nikon) and orca flash 4.0 camera (Hamamatsu). High-speed confocal imaging was performed with a Dragonfly 500 series spinning disk microscope (Andor, Oxford instruments) using a Zyla 4.2 Plus sCMos camera (Andor) and a Lamba CFI-Plan Apochromat 60x 1.4 NA oil immersion objective (Nikon). Three-dimensional data were processed using Fusion software (Andor), and Fiji ImageJ Version 2.1.0. Deconvolution was performed using Huygens Professional 16.10.

### Mitochondria size analysis using Mitograph V2.0

Mitograph image processing was performed according to (*8*). In short, images were processed at resolution level 1 (2x2 binning), corresponding to pixel widths in of 0.2(x) x 0.2(y) x 0.2(z) μm. Data were deconvolved using Huygens professional version 220.10 (SVI) and processed in Fiji version 2.1.0 using the GenFramesMaxProjs.ijm to generate a z-stack and CropCells.ijm supplied from the MitoGraph software to create single cell cropped images. MitoGraph software was run in Terminal with the following parameters (*8*):./MitoGraph -xy 0.2 -z0.2 -adaptive 1 – path ∼/Desktop/Experiments/cells.

### Cell, vacuole and cytosol volume measurements

Cell volumes were determined by fluorescence imaging of Om45-GFP. Signal intensities were increased until cellular outlines were visible. Cell outlines were traced using the ImageJ/Fiji spherical tool to measure long and short axes, and cell volumes were calculated as oblate spheroid shapes. The accuracy of this approach was verified by comparing the data tomeasuring cellular volumes by brightfield microscopy as described in (*6*). Individual vacuoles in yeast show sphericity of 0.97 (*22*). Thus, individual vacuoles for each cell were visualized after vital FM4-64 staining by fluorescence imaging, midsection were traced using the ImageJ/Fiji spherical tool to determine long and short axes, and vacuole volumes were calculated as oblate spheroid shapes. Cytosol volume was approximated as non-vacuolar cell volume; determined vacuole volumes were subtracted from corresponding cell volumes for each individual cell.

### Recombination induced tag exchange (RITE) for monitoring Om45 synthesis

1 OD600-unit of cells were shifted to SD-N media and incubated at 30°C and 180 rpm. After 6 h, β-estradiol was added to a final concentration of 1 μM and cells were incubated for an additional 18 h to induced Cre-recombinase-mediated tag exchange. Protein levels of pre-existing Om45-GFP or newly synthesized Om45-RFP were determined before (0 h) and after starvation (24 h) by dual color fluorescence imaging and line diagram analysis of signal intensities in ImageJ/Fiji. Cre recombinase-induced marker switch was assessed for each genotype by plating 300 cells before and after β-estradiol induction at the end of starvation (24 h) on YPD or YPD + hygromycin plates. Cre recombinase-mediated maker switching was unaffected by tested genotypes.

### Analysis of oxygen consumption rates

XFe96 cell culture microplates (Agilent) were coated with 60 µl 0.1% (w/v) poly-L-lysine (Sigma) for 1 h, washed twice with 100 µl PBS, and stored at room temperature overnight. 0.25 or 0.1 OD_600_-units of cells were harvested and added to the precoated plate with SD-N or SDC, respectively, and incubated for 1 hour at 30°C. Samples were analyzed in a Seahorse XF analyzer (Agilent). The sensor cartridge was loaded with 20 µl FCCP (250 μM; Tebu-bio) in port A, and 20 µl antimycin A (250 μM; Sigma) in port B, to a final working concentration of 25 μM each. The measuring protocol included 3 initial mixing steps, and 3 measurements for the basal levels. Port A and B injection protocols were 3 mixing steps, incubation for 5 min, and three measurements. After the assay, the optical densities were measured by transferring the cultures to a Nunclon Delta Surface 96 well plate in a Varioskan Flash plate reader (ThermoFischer). The data was analyzed and normalized using the Agilent WAVE software version 2.6.1.

### Analysis of cellular reactive oxygen species (ROS)

To measure ROS production, 10^7^ cells were resuspended in media containing 5 μM DHE (Molecular Probes) and incubated for 15 min at 30°C. Using the PE-Cy5 channel of the BD FACS Canto II flow cytometer with 488 nm excitation and ≥670 nm emission settings, the mean fluorescence intensities (MFIs) of 10,000 events per sample were determined.

### Whole cell proteomics

For SILAC during starvation, a concentration of 30 mg/l L-lysine-^13^C_6_,^15^N_2_ hydrochloride (Lys^8^, Sigma, 608041) in SD-N media was determined to show identical autophagic turnover of 2GFP-Atg8 or mitophagic turnover of Om45-GFP compared with SD-N media without lysine addition. For proteomics analysis, cells were grown overnight to 1 OD_600_/ml in SDC, washed three times and resuspended in SD-N. For SILAC, cells were pelleted after 6 h or 12 h and resuspended in SD-N containing 30 mg/l L-lysine-^13^C_6_,^15^N_2_ hydrochloride (Lys^8^, Sigma, 608041) and incubated for indicated time. 0.2 OD_600_-units of cells were harvested and snap frozen in liquid nitrogen.

For cell lysis and protein pellet isolation, a pre-cooled 5 mm stainless steel ball was added to each sample and the cells were pulverized for 1 minute at 25 Hz using a ball mill grinder, equipped with a 48-sample holder (Tissue Lyser 2, Qiagen). The samples were treated with 1 ml pre-cooled (-20°C) mixture of methyl tert-butyl ether (MTBE, Sigma 306975), Optima LC-MS grade methanol (A456-212) and LC-MS Ultra grade water (Honeywell, 14263-1L) in a 50:30:20 ratio. Sample were vortexed immediately until cell pellets were homogenously resuspended. Samples were incubated on an orbital mixer at 4°C for 30 min at 1500 rpm, centrifuged for 10 min at 21100 x g at 4°C and supernatants were removed. For protein digestion, pellets were resuspended in 30 μl 8 M Urea (Sigma). After addition of 0.4 μl 250 mM TCEP and 0.55 μl 1 M CAA, samples were incubated at room temperature for 1 h. 0.5 μl Lys-C (1 μg/μl Lys-C Endoproteinase MS Grade, 90051, Life Technologies) was added and samples were incubated for 2 h at room temperature. Then, 90 μl 50 mM ammonium bicarbonate (VWR) were added. For Non-SILAC samples, 1 µl trypsin (1 µg/µl Trypsin Gold, Mass Spectrometry Grade (Promega)) was added and samples were incubated at 37°C overnight. SILAC samples were incubated at room temperature overnight without trypsin treatment. To stop digestion, the pH was lowered by adding formic acid (FA, VWR) to 1%. Peptides were cleaned and isolated using 30 μg C18-SD StageTips. StageTips were pre-washed in 200 μl methanol, 200 μl 40% (v/v) acetonitrile (ACN, VWR)/0.1% (v/v) FA and 200 μl 0.1% (v/v) FA. Samples were added to the StageTip columns and span for 2 minutes, washed twice with 200 μl 0.1% (v/v) FA. Peptides were eluted into a fresh Eppendorf tube with 40% (v/v) ACN/0.1% (v/v) FA, centrifugation at 300 x g for 4 minutes. Samples were aliquoted into 4 μg/μl using a NanoDrop (Thermofischer), and dried in a SpeedVac and stored at -20°C. 4 mg eluted peptides were dried and reconstituted in 9 µl of 0.1 M TEAB. Labeling with Tandem Mass Tags (TMT, TMT10plex^TM^ or TMTpro™ 16plex, Thermo Fisher Scientific cat. No 90110) was carried out according to manufacturer’s instruction with the following changes: 0.8 mg TMT10plex^TM^ reagent was re-suspended in 70 µl anhydrous ACN; 0.5 mg TMTpro™ 16plex reagent was re-suspended in 33 µl anhydrous ACN. 7 µl TMT reagent in ACN were added to 9 µL peptide/0.1M TEAB. Final ACN concentration was 43.75% and peptides to TMT reagent ratio was 1:20. After 60 min incubation, the reaction was quenched with 2 µl 5% hydroxylamine. Labelled peptides were pooled, dried, re-suspended in 200 µl 0.1% formic acid (FA), split into two equal parts, and desalted using home-made STAGE tips (*27*). One of two parts was fractionated on a 1 mm x 150 mm ACQUITY column, packed with 130 Å, 1.7 µm C18 particles (Waters cat. no SKU: 186006935), using an Ultimate 3000 UHPLC (Thermo Fisher Scientific). Peptides were separated using a 96 min segmented gradient with buffer A (5% ACN, 10mM ammonium bicarbonate (ABC)) and 1% to 50% buffer B (80% ACN, 10mM ABC) for 85 min and from 50% to 95% buffer B for 11 min, at a flow of 30 µl/min. Fractions were collected every three minutes, and fractions were pooled in two passes (1 + 17, 2 + 18 … etc.) and dried in a vacuum centrifuge (Eppendorf). Dried fractions were re-suspended in 0.1% formic acid (FA) and separated on a 50 cm, 75 µm Acclaim PepMap column (Product No. 164942 Thermo Fisher Scientific) using an EASY-nLC1200 (Thermo Fisher Scientific). The analytical column was operated at 50°C. The separation was performed using a 90 min linear gradient with buffer A (0.1% FA) and 6% to 31% buffer B (0.1% FA, 80% CAN). Eluting peptides were analyzed on an Orbitrap Lumos Tribrid mass spectrometer (Thermo Fisher Scientific) equipped with a FAIMS device (Thermo Fisher Scientific). The FAIMS device was operated at -50V and -70V compensation voltages. Mass spectrometric data were acquired in a data-dependent manner with a top speed method. For MS1, the mass range was set to 350−1500 m/z and resolution to 60 K. Maximum injection time was 50 ms and the AGC target to 4e5. Peptides were fragmented using collision-induced dissociation; collision energy was set to 35%. Peptide fragment MS2 spectra were acquired in the ion trap with a maximum injection time of 50 ms and “Turbo” scan rate, using an AGC target of 1e4. The ten most abundant peaks were subjected to Synchronous Precursor Selection and fragmented using higher-energy collisional dissociation; collusion energy was set to 65%. The resulting MS3 spectra were acquired in the Orbitrap at a resolution of 50 K. Raw files were split based on the FAIMS compensation voltage using FreeStyle (Thermo Fisher Scientific).

Raw data were analyzed using MaxQuant (*28*) version 1.6.10.43 for the TMT10plex^TM^ data or version 1.6.17.0 for the TMTpro™ 16plex data, using the integrated Andromeda search engine (*29*). Peptide fragmentation spectra were searched against the canonical sequences of the yeast reference proteome (UniProt proteome ID UP000002311, downloaded October 2018). Methionine oxidation, protein N-terminal acetylation, and Lys^8^ were set as variable modifications; cysteine carbamidomethylation was set as fixed modification. The digestion parameters were set to “specific” and “LysC/P,” The minimum number of peptides and razor peptides for protein identification was 1; the minimum number of unique peptides was 0. Protein identification was performed at a peptide spectrum matches and protein false discovery rate of 0.01. The “second peptide” option was on. Quantification mode was set to “Reporter ion MS3”. The isotope purity correction factors, provided by the manufacturer, were included in the analysis. Data wrangling and exploratory data analysis were performed using the tidyverse package (Wickham, 2019, DOI: 10.21105/joss.01686) in R (R Core Team, 2017). Protein TMT reporter intensity values for the light and heavy SILAC channel were calculated separately from the peptide level data in modificationSpecificPeptides. Differential expression analysis was performed using limma (*30*). To determine the relative incorporation rate (rel. IR), TMT intensities were normalized to the sum of all TMT intensities within one TMT batch. Based on this fraction, MS1 intensities were estimated for each sample resulting in a MS1 intensity for heavy and light for each peptide. MS1 intensity data were used to calculate specific heavy/light ratios (H/L) for each peptide.

Mitochondrial proteins were categorized according to the MitoMAP dataset (REF). To analyze the relative protein composition of mitochondrial proteomes as shown in **figure 2B**, the median abundance for mitochondria proteins was determined for each strain analyzed. Relative differences from the median were calculated for each protein and the resulting differential matrices were compared between genotypes and expressed as Pearson’s coefficients.

### Lipidomics and metabolomics analyses

Cells were grown overnight to 1 OD_600_/ml in SDC and then washed three times and resuspended in SD-N. For indicated timepoints, 1 OD_600_-unit of cells was snap frozen in liquid nitrogen. For α-D-^13^C_6_-glucose tracing, cells were pelleted and resuspended in SD-N containing 2% (w/v) ^13^C_6_:^12^C_6_ α-D-glucose (1:1) at indicated timepoints, incubated for 30 min, and then harvested and snap frozen. The samples were treated with 1 ml pre-cooled (-20°C) mixture of methyl tert-butyl ether (MTBE, Sigma 306975), Optima LC-MS grade methanol (A456-212) and LC-MS Ultra grade water (Honeywell, 14263-1L) in 50:30:20 volume ratio. Additionally, internal standards of 10 µl U-^13^C^15^N amino acids (2.5 mM), 10 µl ^13^C_10_ ATP (1 mg/ml), 20 µl citric acid D4 (100 µg/ml), and 20 µl EquiSPLASH™ LIPIDOMIX. After addition of the extraction buffer, each sample was immediately vortexed until cell pellets were homogenously resuspended. Samples were then incubated on an orbital mixer at 4°C for 30 minutes and at 1500 rpm. Samples were centrifuged for 10 min at 21100 x g at 4°C and supernatants were transferred to a new tube, leaving the protein pellet. 200 µl MTBE and 150 µl LC-MS grade H_2_O was added, and incubated for 10 min at 15°C. Samples were then centrifuged for 5 min at 15°C at 16,000 x g to obtain phase separation. 650 µl of the upper phase, which contains lipids, were transferred to a new tube. The lower phase, which contains metabolites, was transferred to a new tube. Samples were dried in a Speed Vac at 20°C 1000 rpm for 4 hours, and then frozen at -80°C

#### Liquid Chromatography-High Resolution Mass Spectrometry (LC-HRMS) analysis of lipids

Dried lipid samples were resuspended in 250 µl UPLC grade acetonitrile: isopropanol (70:30 [v:v]) mixture, vortexed and incubated for 10 min at 10°C. Samples were centrifuged for 5 min at 10.000 x g, and cleared supernatants were transferred to 2 ml glass vials with 200 µl glass inserts (Chromatography Zubehör Trott, Germany). Glass inserts were placed in a Acquity iClass UPLC (Waters) sample manager at 6°C. The UPLC was connected to a Tribrid Orbitrap HRMS and equipped with a heated ESI (HESI) source (ID-X, Thermo Fischer Scientific). For each lipid sample, 2 µl material were injected onto a 100 x 2.1 mm BEH C8 UPLC column, which was packed with 1.7 µm particles (Waters). The flow rate of the UPLC was set to 400 µl/min. The buffer system included buffer A (10 mM ammonium acetate, 0.1% acetic acid in UPLC-grade water) and buffer B (10 mM ammonium acetate, 0.1% acetic acid in UPLC-grade acetonitrile/isopropanol 7:3 [v/v]). The gradient for the UPLC was as follows: 0-1 minu 45% A, 1-4 min 45-25% A, 4-12 min 25-11% A, 12-15 min 11-1% A, 15-18 min 1% A, 18-18.1 min 1-45% A and 18.1-22 min re-equilibrating at 45% A. This results in a total runtime of 22 min per sample. The ID-X mass spectrometer was operating either for the first injection in positive ionization mode, or for the second injection in negative ionization mode. In either case, the analyzed mass range was between m/z 150-1500. The resolution was set to 120.000 resulting in approximately 4 scans per second. The RF lens was set to 60%, and the AGC target was set to the value 500% (∼2E6 ions). 100 ms was set to the maximal ion time and the HESI source was operating with a spray voltage of 3.5 kV in positive ionization mode, additionally 3.2 kV were applied in negative ionization mode. The capillary temperature was set at a value of 250°C, the sheath gas flow at 60 arbitrary units (AU), the auxiliary gas flow at 20 AU and the sweep gas flow was set to 2 AU at a value of 350°C. The run-order for all samples was randomised and the data were analysed using a targeted approach employing the quan module of the TraceFinder 4.1 software (Thermo Fischer Scientific) in combination with a sample-specific in-house generated compound database.

#### Anion-Exchange Chromatography Mass Spectrometry (AEX-MS) for anionic metabolite analysis

Extracted metabolites were resuspended in 150 µl Optima LC/MS grade water (Thermo Fisher Scientific) of which 100 µl was transferred to p*olypropylene* autosampler vials (Chromatography Accessories Trott, Germany). The samples were analyzed using a Dionex ionchromatography system (Integrion, Thermo Fisher Scientific) as described previously in (*31*). In brief, 5 µl polar metabolite extract were injected in push partial mode using an overfill factor of 1 onto a Dionex IonPac AS11-HC column (2 mm × 250 mm, 4 μm particle size, Thermo Fisher Scientific), additionally this was equipped with a Dionex IonPac AG11-HC guard column (2 mm × 50 mm, 4 μm, Thermo Fisher Scientific). The column temperature was held at 30°C and the auto sampler was set to a temperature of 6°C. To generate a potassium hydroxide gradient, potassium hydroxide cartridge was used (Eluent Generator, Thermo Scientific), in addition to deionized water. The metabolite separation was carried at a flow rate of 380 µl/min, applying the following gradient conditions: 0-3 min, 10 mM KOH; 3-12 min, 10−50 mM KOH; 12-19 min, 50-100 mM KOH, 19-21 min, 100 mM KOH, 21-22 min, 100-10 mM KOH. The column was finally re-equilibrated at 10 mM for 8 min.

For the analysis of metabolic pool sizes, the compounds were eluted and detected in negative ion mode using the full scan measurements in the mass range m/z 50 – 750 on a Q-Exactive HF high resolution MS (Thermo Fisher Scientific). The heated electrospray ionization (ESI) source settings of the mass spectrometer were: Spray at a voltage of 3.2 kV, the capillary temperature was set to a temerpature of 300°C, sheath gas flow set to 60 AU and aux gas flow set to 20 AU, and at a temperature of 300°C. The S-lens was set to a value of 60. The mass spectrometric data analysis of each compound was performed using the TraceFinder software (Version 4.1, Thermo Fisher Scientific). The identification of each compound was validated by authentic compound references, which were measured in alongside the experimental sample set. For data analysis, the deprotonated [M-H^+^]^-^ monoisotopic [M0] mass peak area of each compound was extracted, and then integrated using a mass accuracy <5 ppm and a retention time (RT) tolerance of <0.05 minutes as a threshold, and this was compared to the measured reference compounds, independently. Pool sizes were normalized to the added internal standards from the extraction buffer, followed by a normalization to the optical density (OD) of the sample.

#### Untargeted liquid chromatography-high-resolution mass spectrometry-based (LC-HRS-MS) analysis of amine-containing metabolites

For the LC-HRMS analysis of amine-containing compounds, the protocol was performed using an adapted derivatization benzoylchlorid-based method (Wong *et al.*, 2016). The polar fraction of each metabolite sample extract was resuspended in 120 µl LC-MS-grade water (Optima-Grade, Thermo Fisher Scientific). 30 µl cleared supernatant were mixed in a critical clean autosampler vial with the addition of a 200 µl glass insert (Chromatography Accessories Trott, Germany). The extract was mixed with 15 µl 100 mM sodium carbonate (Sigma), and 15 µl 2% [v/v] benzoylchloride (Sigma) in acetonitrile (Optima-Grade, Thermo Fisher Scientific). Samples were vortexed and kept at 20°C until analysis. For the analysis, 1 µl derivatized sample was directly injected onto a 100 x 2.1 mm HSS T3 UPLC column (Waters). The flow rate of 400 µl/min was set using a buffer system consisted of buffer A (10 mM ammonium formate (Sigma), 0.15% formic acid (Sigma) in Milli-Q water (Millipore)) and buffer B (acetonitrile, Optima-grade, Fisher-Scientific). The column temperature was set to 40°C, while the LC gradient was: 0% B for 0 – 4.1 min; 0-15% B for 4.1 – 4.5 min; 15-17% B for 4.5-11 min; 17-55% B for 11 –11.5 min, 55-70% B for 11.5 – 13 min; 70-100% B for 13 – 14 min; 100% B for 14 -14.1 min; 100-0% B for 14.1-19 min; 0% B. The mass spectrometer operated in positive ionization mode recording the mass range m/z 100-1000. The settings for the heated ESI source of the mass spectrometer were: Spray voltage set to 3.5 kV, capillary temperature of 300°C, sheath gas flow of 60 AU and aux gas flow of 20 AU at a temperature of 340°C. The S-lens was set to a value of 60 AU. Data analysis was performed using the TraceFinder software (Thermo Fisher Scientific). Reference compounds were used to identity each compound. Peak areas of [M + nBz + H]^+^ ions were extracted using a mass accuracy (<5 ppm) and a retention time tolerance of <0.05 min.

## Notes

### Competing Interest Statement

The authors have declared no competing interest.

