## Supplementary figures and images for "Homeostatic mitophagy scales mitochondrial networks"

### Supplementary Figure S1

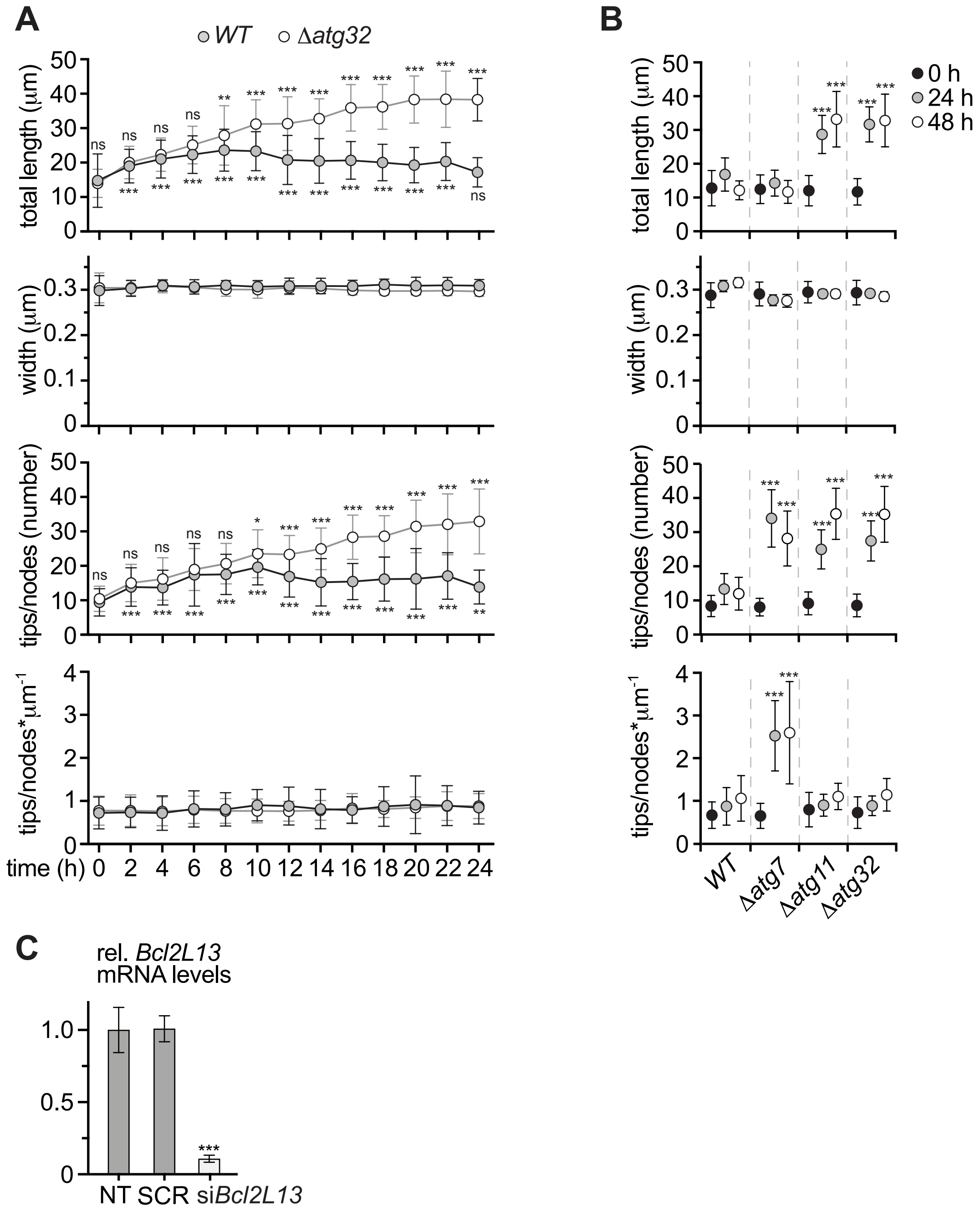


**Supplementary Figure 1**

### Supplementary Figure S2

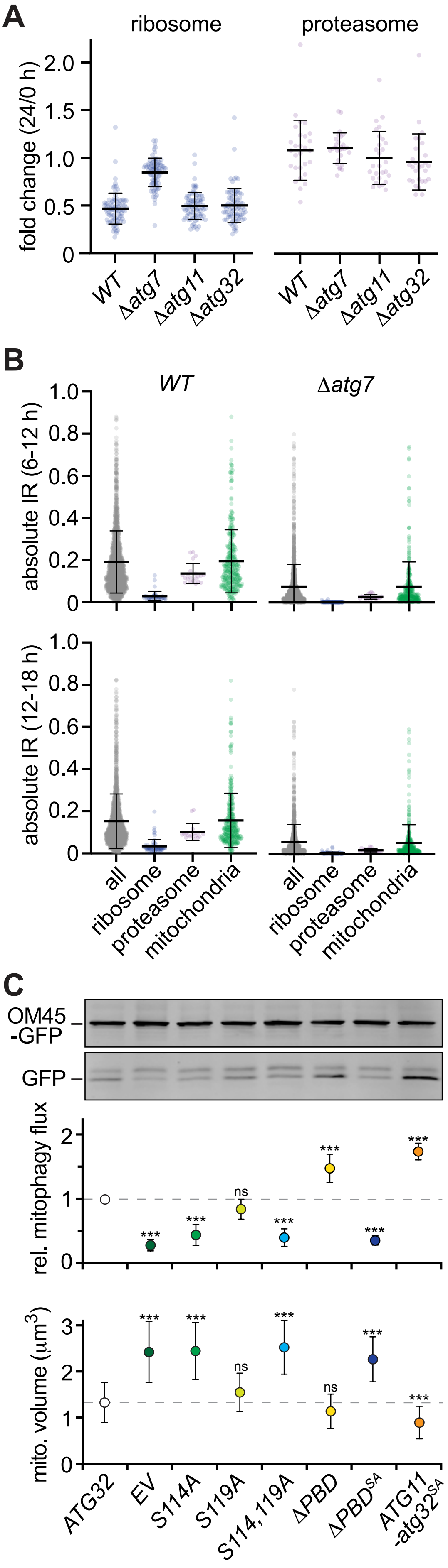


**Supplementary Figure 2**

### Supplementary Figure S3

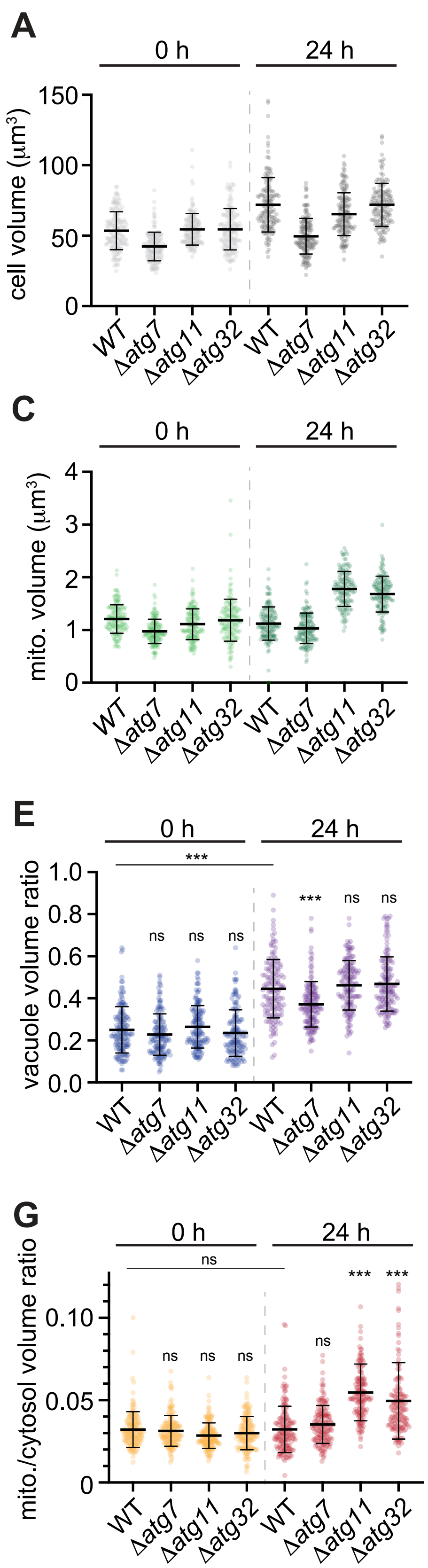


**Supplementary Figure 3**

### Supplementary Figure S4

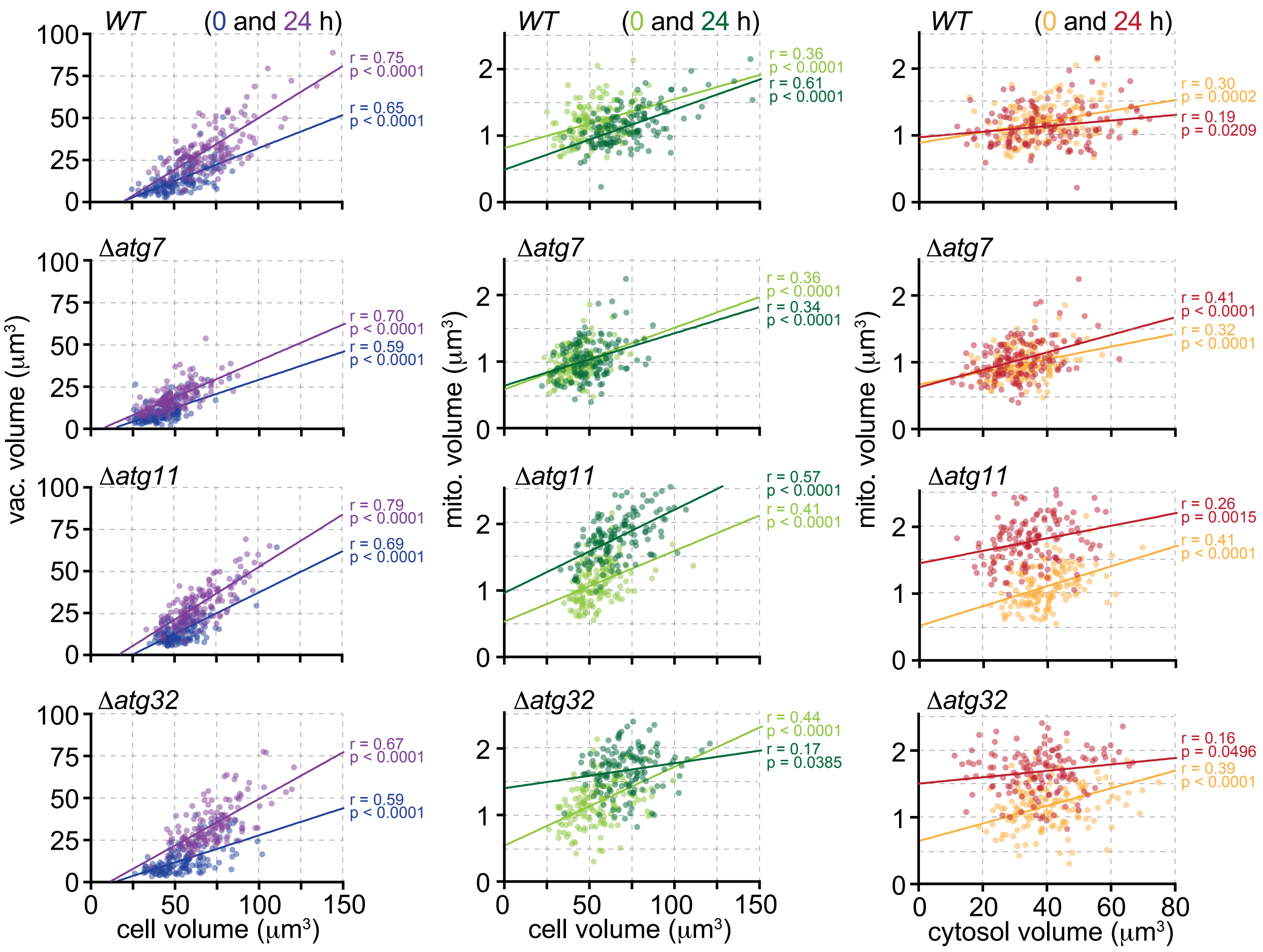


**Supplementary Figure 4**

### Supplementary Figure S5

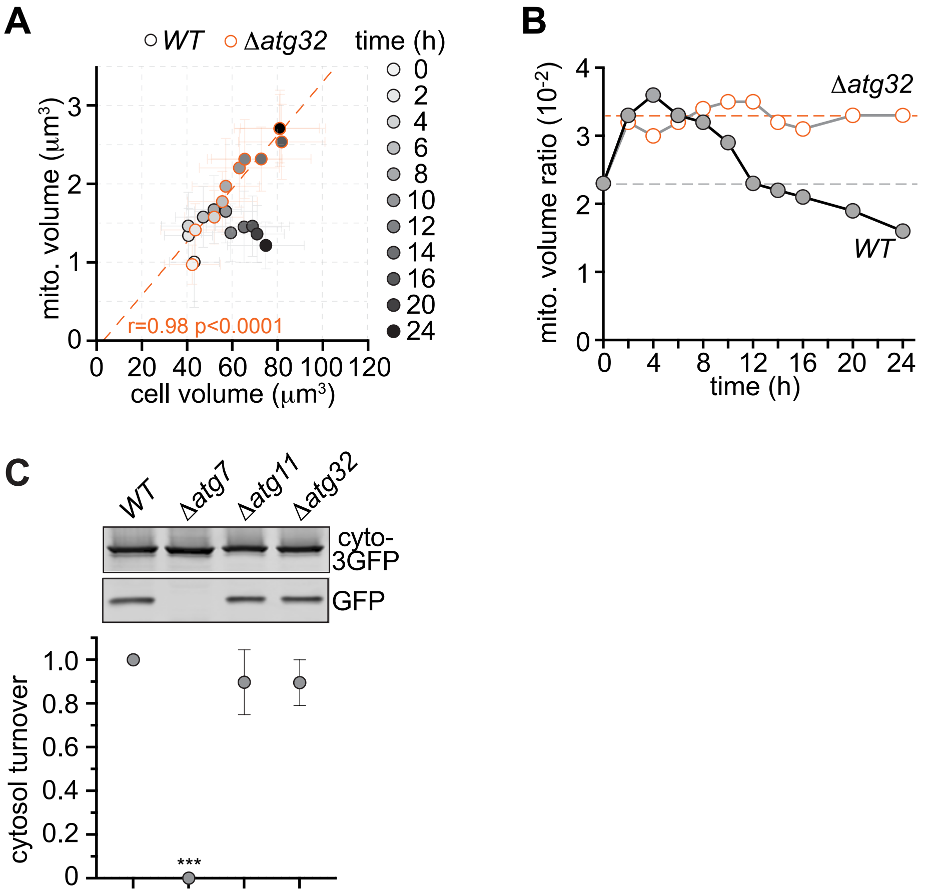


**Supplementary Figure 5**

### Supplementary Figure S6

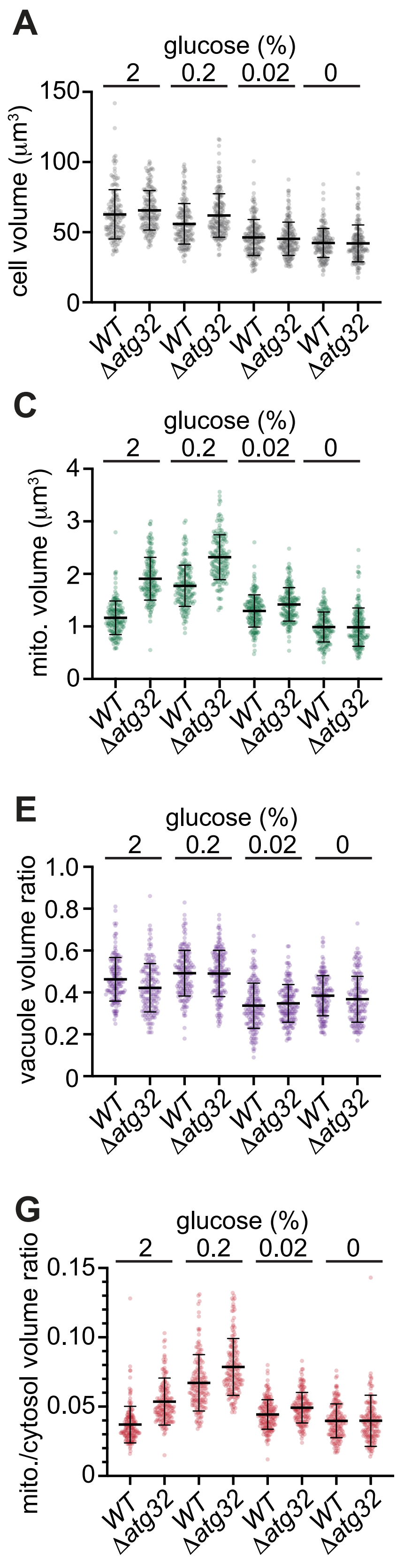


**Supplementary Figure 6**
